# Nitrogen-fixing *Klebsiella variicola* in feces from herbivorous tortoises

**DOI:** 10.1101/666818

**Authors:** Diana Montes-Grajales, Berenice Jiménez, Marco A. Rogel, Alejandro Alagón, Nuria Esturau-Escofet, Baldomero Esquivel, Julio Martínez-Romero, Esperanza Martínez-Romero

## Abstract

Animals feeding on plants (herbivorous) may have nutritional deficiencies and use bacterial nitrogen fixation in guts to compensate unbalanced diets with high carbon and low nitrogen. Using the acetylene reduction assay we searched for nitrogen fixation in the feces from several herbivorous animals in captivity. We detected acetylene reduction in feces from two African spurred tortoises, *Centrochelys sulcata* and in feces from six *Gopherus berlandieri* tortoises and isolated nitrogen-fixing klebsiellas from them. Additionally, we performed a gut metagenomic study with Illumina sequencing from a healthy Mexican *G. berlandieri* tortoise, and the *nif* genes identified in the feces microbiome matched those from *Klebsiella variicola*. Fecal bacterial composition from tortoises was similar to that reported from other reptilian guts.

## Introduction

Nitrogen fixation is a unique biochemical process carried out by nitrogenases encoded by *nif* genes, which are found in only a few prokaryotes (1). Nitrogen fixation is energetically expensive and nitrogen-fixing bacteria are in many cases minor components of the microbial community that nevertheless provide a valuable ecological service (2). Nitrogen-fixing bacteria are called diazotrophs (3).

When associated with nitrogen-fixing bacteria, insects or plants may inhabit in nitrogen poor conditions. Termites contain nitrogen-fixing bacteria in their guts that allow them to grow in wood, similarly wood eating beetles contain nitrogen-fixing bacteria (4). Nitrogen fixation by *Klebsiella variicola* occurs in the fungal garden of ants (5). A novel betaproteobacterium capable of fixing nitrogen is a recently described symbiont that is transmitted by eggs of carmine cochineals which feed on cacti as the *Gopherus berlandieri* tortoise does (6). Cacti have high sugar content and are nitrogen poor, conditions that would be favorable for nitrogen fixation in animals that use cacti for food, such as the carmine cochineal or the *G. berlandieri* tortoise.

*G. berlandieri*, is distributed from North Mexico (states of Coahuila, Nuevo Leon and Tamaulipas) to South-central Texas in United States (7). This species generally maintains restricted mobility by living for prolonged times in the same burrow (8) and its diet is mainly composed of grasses, weeds and cacti. On the other hand, the African spurred tortoise, *Centrochelys sulcata*, which is also vegetarian, lives in West Africa (9).

Metagenomic analyses have contributed to gain a better insight on taxonomic composition and functions of the gut microbiota of various species, including non-model organisms, such as sea turtles (10). Gut microbiota may confer adaptability to herbivorous tortoises such as *G. berlandieri* and *C. sulcata* naturally exposed to low nitrogen-content diets. The aim of this research was to identify nitrogen-fixing species in the gut microbiota of *G. berlandieri* and *C. sulcata*. In addition we present one *G. berlandieri* fecal metagenome analysis and nuclear magnetic resonance (NMR) spectroscopy-based identification of the exometabolites produced by one of the nitrogen-fixing bacterial isolates from the gut microbiota of *G. berlandieri* and by *K. variicola* F2R9 as reference.

## Materials and methods

### Samples

Six samples -each from a different individual- were obtained from *G. berlandieri* turtles kindly supplied by Ezer Yniestra with permission of SEMARNAT. They were fed with a vegetarian diet based mainly on cactus cladodes, grass, clover, dandelion and lettuce and received a dietary supplement based on vegetables once a week. They were dewormed annually and never received antibiotics. Another tortoise species with similar habits and life conditions is the African turtle *C. sulcata*. A *C.sulcata* stool sample was provided by Zoofari local zoo and used to detect acetylene reduction, additional samples were from Dr. Alejandro Alagón’s *C. sulcata* tortoise, which has been also maintained under a vegetarian diet.

All stool samples were collected using sterile falcon tubes and stored at −20°C. The unique sample used for metagenomic DNA extraction and sequencing, consisted of feces from a female adult *G. berlandieri* tortoise from the herpetarium of the National Autonomous University of Mexico Faculty of Sciences, located in Mexico City (Mexico). This followed an herbivorous diet and did not take antibiotics at least the year before the sample was taken.

### Nitrogen-fixing bacteria isolation and acetylene reduction assays

Samples of feces of six individuals of *G. berlandieri* and one of *C. sulcata* were employed for nitrogen-fixing bacteria isolation and molecular identification, as well as food samples of *G. berlandieri* consisting on a mixture of 85% of cactus (*nopal*) and 15% lettuce. Glass vials of 10 ml were filled with 5 ml of soft agar nitrogen free medium (Na_2_HPO_4_·12H_2_O 0.22% w/v; NaH_2_PO_4_·H_2_O 0.0425% w/v; MgSO_4_·7H_2_O 0.04375% w/v; C_12_H_22_O_11_ 0.1%; Fahraeus traces 0.1% v/v and agar 0.22%). Six replicates of each sample were inoculated and incubated for 2 days into the medium (3 replicates at 30°C, and the others at 37°C). After that cotton plugs where changed for hermetic rubber caps, 0.6 ml of acetylene were injected in the flasks and incubated for 3 days. A volume of 0.4 ml of air from each vial was used for ethylene and acetylene detection by gas chromatography (GC). In addition, acetylene reduction activity of the stools samples and isolates were tested using different carbon sources, such as sucrose, glucose and D-gluconic acid sodium salt.

LB-agar petri plates were inoculated with bacterial growth from the vials that presented acetylene reduction and incubated for 24h at 30°C or 37°C. Bacteria with different phenotypic features from each sample were isolated by the streak plate method and tested for acetylene reduction activity (ARA) by GC, after 24h of growth in glass vials with 5 ml of nitrogen free medium and 24h of incubation with 0.6 μl of acetylene.

A total of 25 bacterial isolates were chosen for DNA extraction and identification of enterobacterial repetitive intergenic consensus (ERIC) PCR patterns. At least one sample of the purified PCR product of each pattern was used for paired end 16S rDNA gene sequencing (Macrogen). Additionally, a BOX-A1R-based repetitive extragenic palindromic-PCR (BOX-PCR) was performed with few selected isolates.

### Phylogenetic analysis of nitrogen-fixing bacterial isolates using 16S rRNA gene sequences and antibiotic susceptibility test

The 16S rDNA gene sequences of the nitrogen-fixing bacterial isolates from *G. berlandieri* and *C. sulcata* stools were submitted to standard nucleotide blast (11). The result of this screening was employed to select the reference strains for the phylogenetic analysis from the NCBI reference sequences database (RefSeq) Alignments were carried out by MAFFT (12); nucleotide substitution model selection was performed with JModelTest (HKY85) (13), which is based on the maximum-likehood principle, and PhyML3.0 software (14) used for phylogenetic analysis with a bootstrap of 100. The phylogenetic tree visualization and editing was carried out in Mega X (15).

Antibiotic susceptibility of the nitrogen-fixing isolated bacteria and reference strains were performed in liquid LB medium with the following antibiotics: carbenicillin 100 μg/ml (Cb100), nalidixic acid 20 μg/ml (Nal20), tetracycline 10 μg/ml (Tc10), chloramphenicol 25 μg/ml (Cm25), kanamycin 50 μg/ml (Km50), neomycin 100 μg/ml (Nm100), spectinomycin 100 μg/ml (Sp100), ampicillin 100 μg/ml (Am100), 20 μg/ml gentamicin (Gm20), streptomycin. 100 μg/ml (Sm100). -Controls with no antibiotics were grown. All samples were kept in glass tubes under shaking (250 rpm), and initial OD600 ~ 0.1, during 24h. Subsequently, he relative bacterial growth was measured by comparison with the respective sample.

### DNA extraction and metagenome sequencing

DNA extraction was performed with the following modifications of the protocol described by Zhang et al. (16). The ethanol wash was performed on ice followed by centrifugation at 4°C; lysis was carried out by using liquid nitrogen, and the incubation with K proteinase was conducted during 4 hours at 45°C. A 1:1 liquid-liquid extraction was done with phenol by shaking gently for 10 minutes and centrifuging at 3000 g for 10 minutes. A second extraction was conducted with phenol:chloroform:isoamyl alcohol 25:24:1 by shaking gently for 10 minutes and centrifuging at 3000 g for 10 minutes. DNA was precipitated using isopropanol 99% overnight at −20°C, followed by centrifugation (30 min, 8000 g) to decant it. The pellet was washed twice with 1ml of 70% ethanol and dried, then resuspended in 30 μl of TE buffer 10/1 and incubated with 10 μl of RNasa (1mg/ml). To avoid the presence of cetrimonium bromide (CTAB), which has a soapy consistency, a third extraction using phenol:chloroform:isoamyl alcohol 25:24:1 was performed by shaking gently for 10 minutes and centrifuging at 16000 g for 10 minutes. DNA was precipitated using cold isopropanol for 3h at −20°C, and a centrifugation step at 16000 rpm for 20 min

Quality was estimated using agarose gel and nanodrop analysis. Raw metagenomic data were obtained by paired-end Illumina HiSeq™ 2500 sequencing technology (Baseclear). Original reads were deposited at the National Center for Biotechnology Information (NCBI).

### *G. berlandieri* metagenome analysis

Illumina paired-end reads were adapter- and quality-trimmed using Trim Galore (17), with a Phred score cutoff of 20. Megahit (18), MetaSpades (19) and IDBA-UD (20) programs where employed to generate three metagenome assemblies. Due to our gene-centric interest, in the search of nitrogen-fixation related genes, high confidence contigs and assembly of a substantial proportion of the dataset was preferred, instead of larger contigs as usual for genome-based approaches (21). Therefore, after quality assessment by QUAST, the more convenient assembly was selected for further analysis using as selection criteria the total length of the assembly and number of contigs (>= 1000 bp).

Taxonomic classification and functional annotation of the trimmed reads were carried out by Kaiju (17), using the non-redundant protein database: bacteria, archaea, viruses, fungi and microbial eukaryotes database (NCBI BLAST nr+euk) and visualized with Krona (22), and HUMAnN2 -The HMP Unified Metabolic Analysis Network- (23), respectively. Gene prediction and annotation was carried out by PROKKA (24). In addition, Blast (Buhler et al., 2007) was used to compare taxonomic predictions with reference genomes, in order to refine the results and avoid errors due to incorrectly annotated sequences. The search of nitrogen-fixing related genes was carried out by Blast (11)with *nif* genes from NCBI, using tblastn with an E-value 1e-50, to obtain a small number of hits with high quality. In addition, blastn was used to identify virulence and related genes in the metagenome with the same e-value, and only those with an identity ≥95% over an alignment length were retained, reference nucleotide sequences were downloaded from EMBL (25).

### Exo-metabolites identification

*K. variicola* type strain F2R9 and the bacterial isolate KGB_5 obtained from *G. berlandieri* were grown on liquid minimal medium and nitrogen free minimal medium, under microaerobic conditions. Culture supernatants were centrifuged and the extracellular medium lyophilized. These freeze-dryed samples were dissolved in 600 μl of D_2_O sodium phosphate buffer 0.123M at pH 7.4 with trimethylsilylpropionic acid (TSP) 1mM as internal standard. An Avance III HD 700 700 MHz NMR spectrometer equipped with a 5-mm z-axis gradient TCI cryogenic probe (Bruker, Fällanden, Switzerland) was used at 298 K and ^1^H frequency of 699.95 MHz. ^1^H NMR Spectra acquisition were carried out by using the standard NOESY-1D pulse sequence (Bruker program *noesypr1d*), which suppresses water preserving the intensity of most of the lasting signals (26). Water signal was irradiated during relaxation delay (RD): 4.0 s and mixing time: 10 ms, 256 scans with spectral width: 14 kHz, 64k data points, acquisition time: 2.3 s and exponential line-broadening factor: 0.3 Hz was applied to the free induction decays (FID) before Fourier transformation. Topspin v 3.5.6 and MestReNova v. 12.0 (MestreLab Research SL.) were used for recording and processing the spectrum (phase and baseline), respectively. TSP chemical shift referenced to 0.000 ppm and Chenomx NMR Suite v. 8.31 (Chenomx Inc.) was used for identification and quantification of the exo-metabolites.

## Results

### Nitrogen-fixing bacteria isolation and acetylene reduction assays

ARA was detected in few fecal samples from herbivorous animals in the Zoofari zoo in Morelos, Mexico. Among them, *C. sulcata* tortoise stool sample was positive for ethylene production in the ARA assay. Further analysis with an extended sampling of tortoise stools showed that acetylene reduction was obtained in all stools tested and in cultures from the isolates from both *G. berlandieri* and *C. sulcate* samples (Table 1, Fig 1). In addition, one of the food samples of *G. berlandieri* was positive for ARA assay, using sucrose as carbon source (12.02 nmol C_2_H_4_/h vial).

**Table 1.** Acetylene reduction activity in *G. berlandieri* and *C. sulcata* stools.

| Sample | Acetylene reduction activity (nmol C<sub>2</sub>H<sub>4</sub>/h vial) |  |  |
| --- | --- | --- | --- |
|  | Glucose | Sucrose | D-gluconic acid sodium salt |
| *G. berlandieri* stool | 167.96±36.23 | 222.72±25.71 | 57.56±20.66 |
| *C. sulcata* stool | 8.31±5.83 | 37.46±5.26 | 0.56±0.19 |
| *K. variicola* F2R9 | 4.23±0.64 | 30.41±0.56 | 7.78±1.37 |
| *K. variicola* VI | 13.77±4.81 | 56.38±7.38 | 12.82±1.83 |

**Fig 1.**
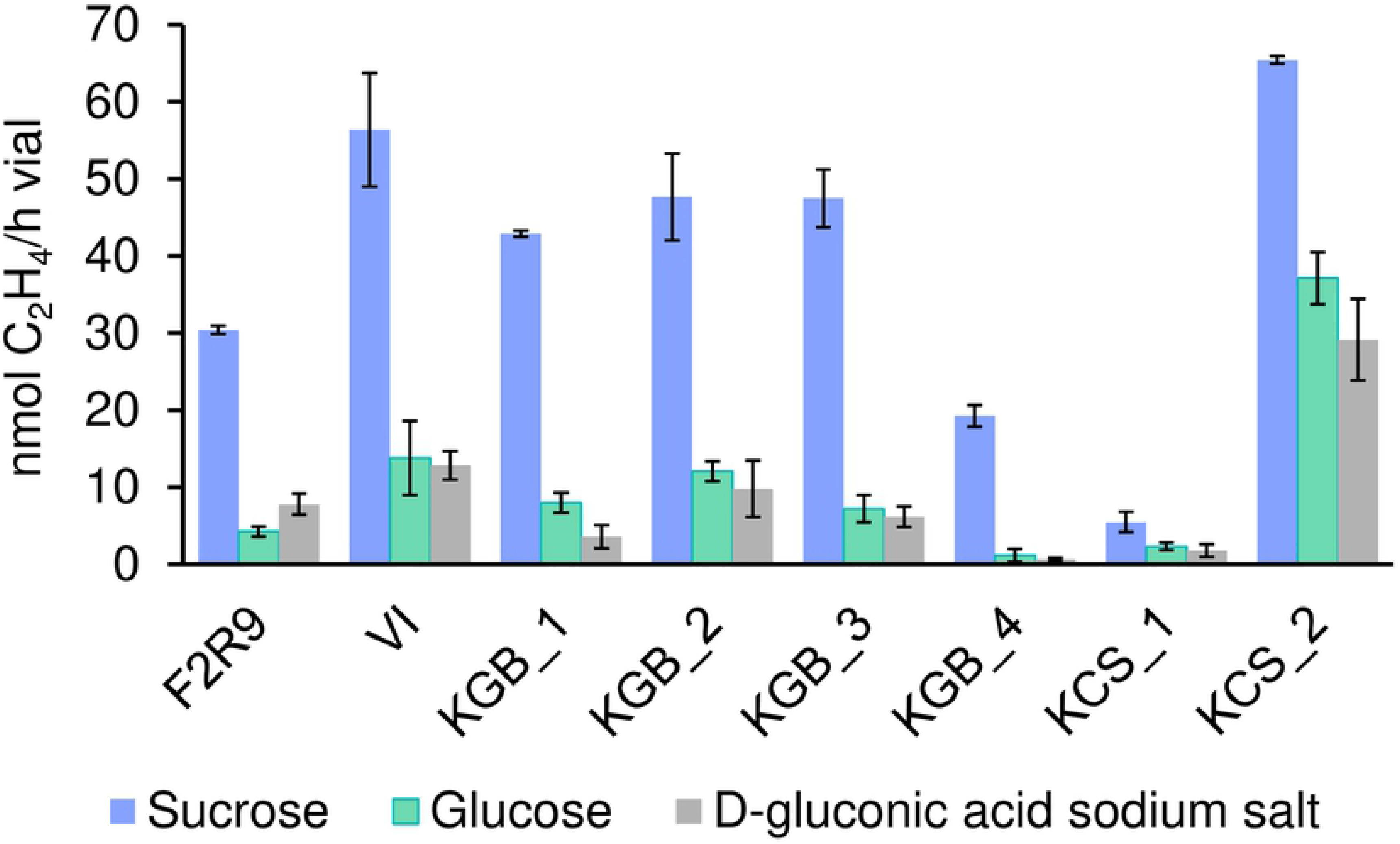
Acetylene reduction activity of bacterial grown in nitrogen-free medium supplemented with different carbon sources.

Nitrogen-fixing strains demonstrated the variation in the amount of acetylene reduced according to the available carbon source (Fig 1). In plant nitrogen-fixing symbioses, carbon supplies have been found to limit nitrogen fixation. Previously, it was reported that *E. coli* mutants affected in gluconate catabolism had a reduced gut colonization (27). Thus we tested if gluconate supported nitrogen fixation in klebsiella and found that it was not the best substrate, seemingly a better supply was sucrose.

A total of 27 bacterial isolates were obtained from *G. berlandieri* (KGB) and *C. sulcata* (KCS) stool samples, 24 and 3, respectively. ARA test showed that 25 of them presented detectable nitrogen fixation by GC, and they exhibited identical colony morphology to *K. variicola* reference strains. According to the ERIC test, six different patterns were observed, and most of them presented the same fingerprints. The patterns of representative strains is shown in Fig 2, fifteen strains from *G. berlandieri* also presented the same pattern as KGB_1 (including KGB_6 shown in Fig 3), three strains from *G. berlandieri* exhibited the same fingerprint as KGB_3 (including KGB5 shown in Fig 3) and one more strain from *C. sulcata* presented KCS1 pattern. Strains KGB2, KCS2 and KGB4 represent each one an unique pattern. In addition, a bacterial isolated from food sample (KFOOD) of *G. berlandieri* also presented acetylene reduction activity, but exhibited a different genomic fingerprint (ERIC pattern) from those in feces. Representative strains of each group were selected for 16s rDNA gene sequencing and acetylene reduction assays.

**Fig 2.**
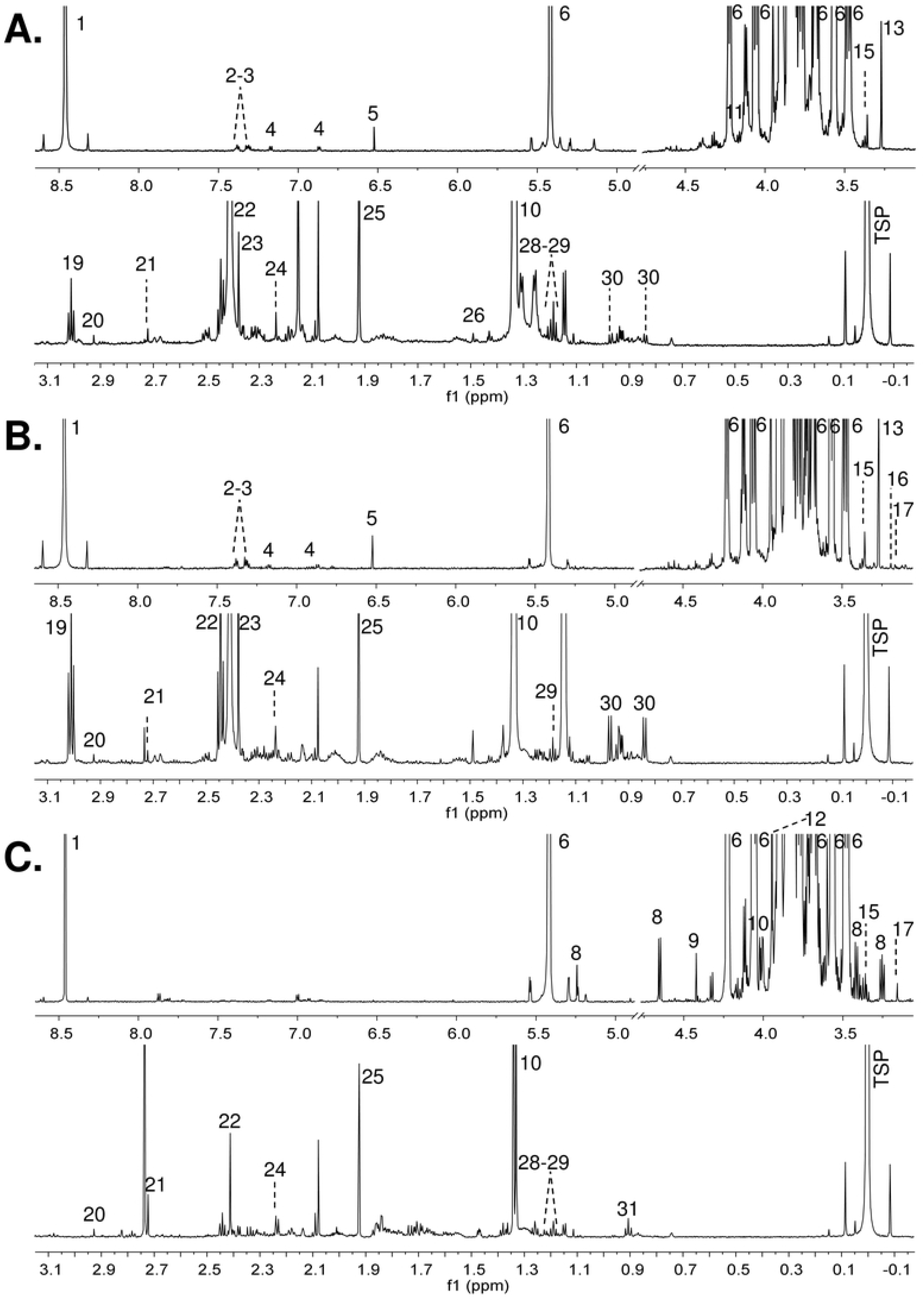
ERIC fingerprints of representative bacterial isolates from *G. berlandieri* and *C. sulcata* stools. M: 1 Kb Plus DNA Ladder, 1: *K. variicola F2R9*, 2: *K. variicola* VI, 3: KGB_1, 4: KGB_2, 5: KCS_1, 6: KGB_3, 7: KGB_4, 8: KCS_2.

**Fig 3.**
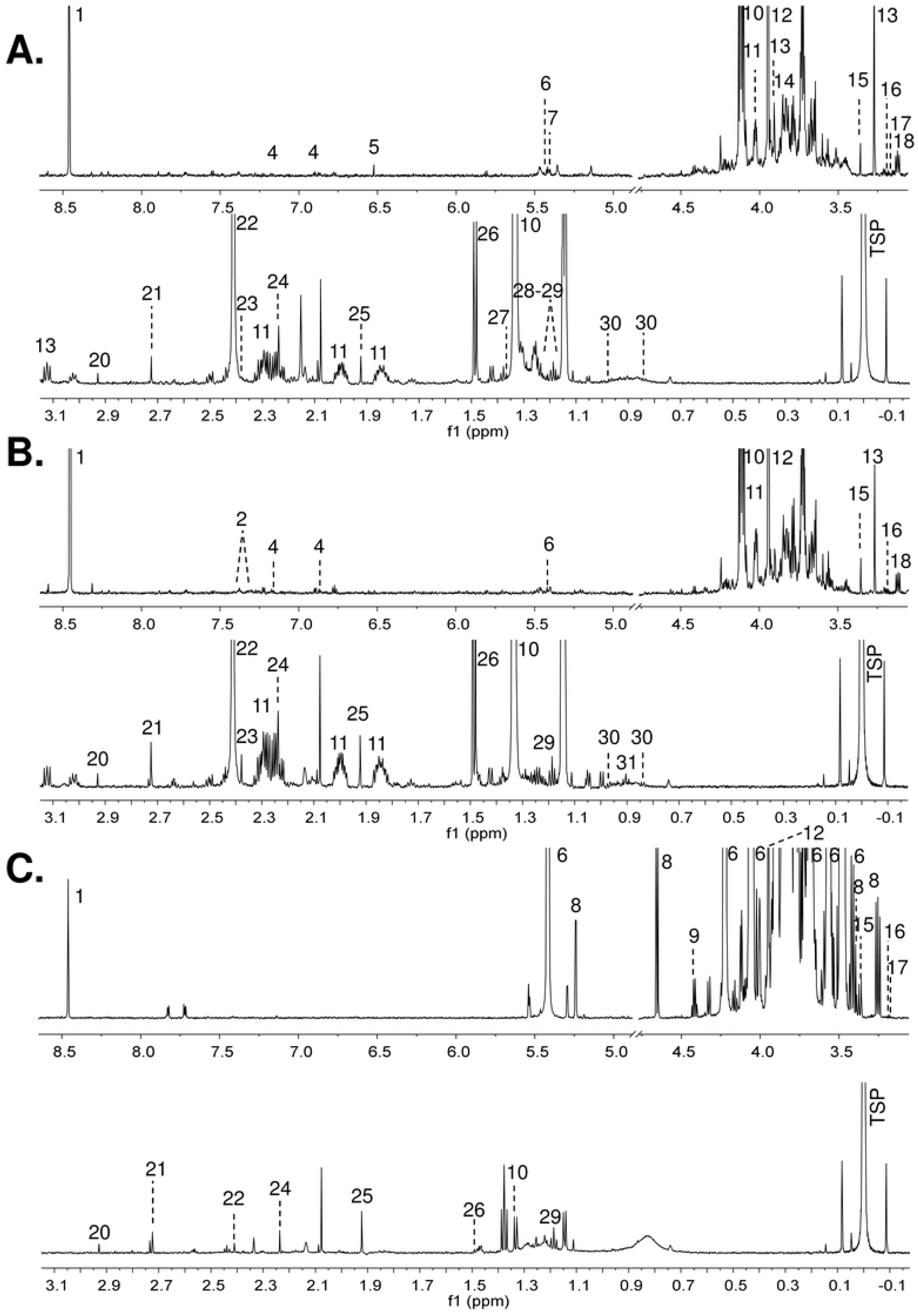
Phylogenetic tree of 16S rRNA gene sequences of representative nitrogen-fixing bacterial isolates from *G. berlandieri (KGB)* and *C. sulcata (KCS)* stools, food of *G. berlandieri* (KFOOD) and reference strains.

### Phylogenetic analysis of nitrogen-fixing bacteria using 16S rRNA gene sequences

The phylogenetic tree obtained indicated that the closest bacteria species to the nitrogen-fixing isolates from *G. berlandieri* and *C. sulcata* is *K. variicola* for most of the strains, and *K. michiganesis* and *K. oxytoca* for one of the isolates from *C. sulcata* and the strain isolated from the food of *G. berlandieri* (Fig 3), which was congruent with fingerprints found by BOX-PCR. The 16S rRNA gene sequence analysis of the non-nitrogen-fixing isolated strains revealed that they were close to *Raoultella ornithinolytica* and *Klebsiella pneumoniae* (not shown).

From a single stool sample, different *Klebsiella* strains were isolated but remarkably from different tortoise samples common strains were obtained.

The results of the antibiotic susceptibility tests shown that the nitrogen-fixing bacterial isolates from *G. berlandieri* and *C. sulcata* in the presence of different antimicrobials, as well as the *K. variicola* reference strains, were resistant to Nm100 and Amp100, and susceptible to Cb100 and Nal20. In general, most of the nitrogen-fixing isolated strains presented the same antibiotic susceptibility pattern as *K. variicola* F2R9.

### DNA extraction and metagenome sequencing

Illumina Hiseq2500 paired-end sequencing of a DNA sample obtained from *G. berlandieri* stool was performed. A total of 84,971,762 read pairs were generated with an average G/C content of 52%. Raw reads for the gut metagenome of *G. berlandieri* were deposited in the NCBI short read archive (SRA) under accession number: SRR8986148.

### *G. berlandieri* metagenome analysis

After trimming, 99.4 % of the original reads were kept for subsequent analysis, corresponding to 84,464,617 cleaned read pairs. The generated MetaSpades, IDBA-UD and MegaHit assembies consisted of 92799, 75327 and 98289 contigs (>= 1000 bp), respectively; accounting for 388140063, 271529393 and 393249585 total length (>= 1000 bp), in each case. Accordingly, Megahit assembly was selected for further identification of nitrogen fixation-related genes in *G. berlandieri* gut microbiota. Due to, Megahit generated the largest assembly with more contigs of size >= 1000 bp.

According to the taxonomic analysis by Kaiju (28), gut microbiota of *G. Berlandieri* is dominated by bacteria (97%) with a prevalence of the Firmicutes fylum (53%), followed by bacteroidetes (10%) and Proteobacteria (9%), in which the most abundant class is Gammaproteobacteria (4%); predominantly shaped by bacteria of the order enterobacteriales (1%), the family Enterobacteriaceae (0.9%) and the genus *Klebsiella* (0.3%) (Fig 4, S1 Table). Archaeas (2%), Eukaryota (0.7%) and viruses (0.07%) were also detected.

**Fig 4.**
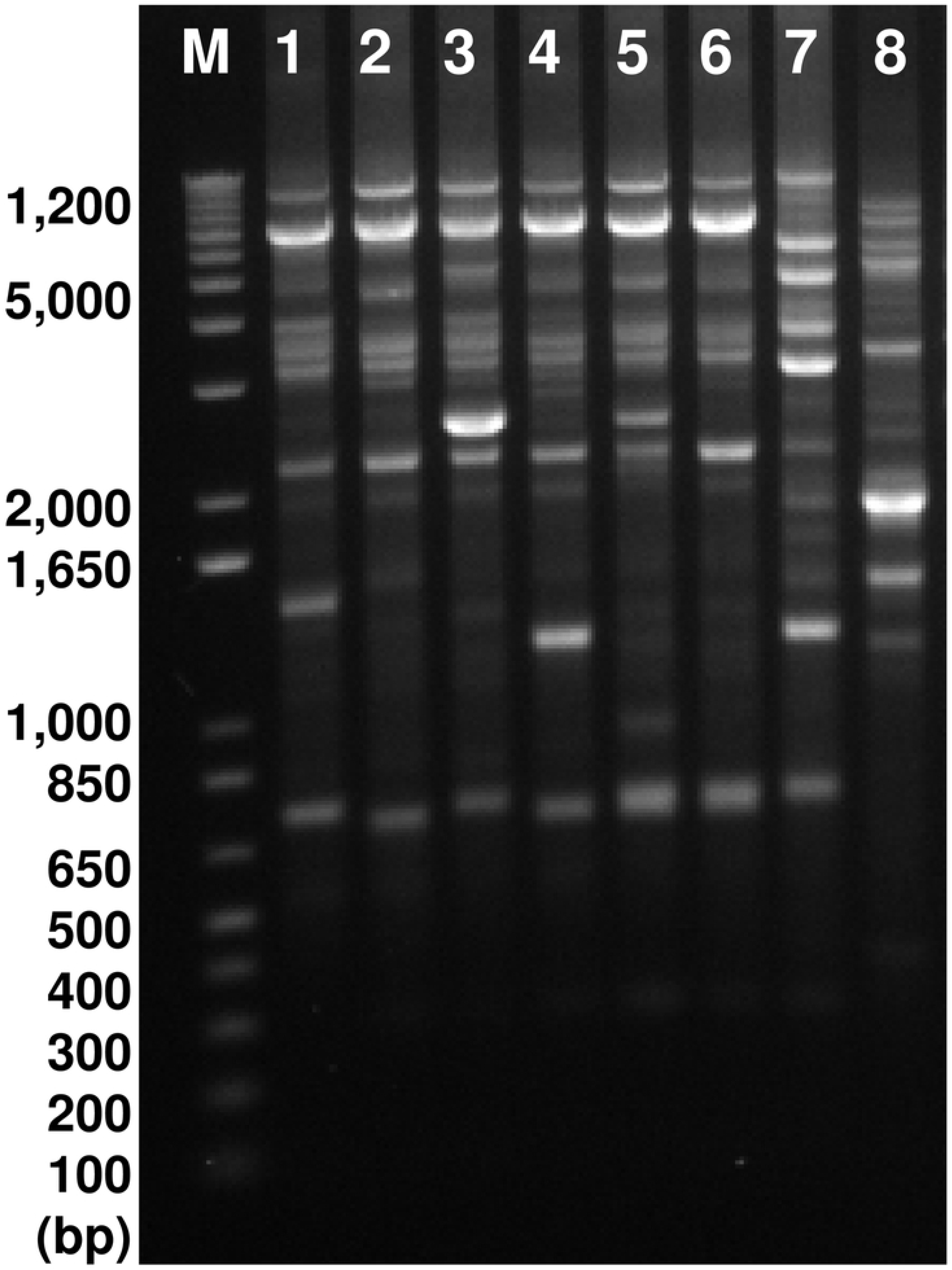
Krona graph representing the taxonomic profile of the gut metagenome of *G. berlandieri*. A. showing the relative abundance of bacteria, archaea, viruses, fungi and microbial eukaryotes; and B. proteobacteria phylum, including *Klebsiella*.

Additional alignment of *K. variicola* F2R9 16s ribosomal RNA gene sequence (NCBI reference sequence: NR_025635.1) against the metagenome was conducted with blastn obtaining an identity 99.7 %, this was the closest species to most of the bacterial isolates in the phylogenetic tree.

Functional annotation, grouping by hierarchical level and normalization of the abundances were carried out using HUMAnN2. A total of 485 pathways were identified in the gut metagenome of *G. berlandieri*, of which 240 were present in klebsiellas (S2 Table). Biosynthetic pathways for most of amino acids such as L-arginine, L-histidine, L-lysine, L-aspartate, L-asparagine, cysteine, L-serine, glycine, L-proline, L-threonine, L-methionine, L-tryptophan and L-alanine were found in the metagenome. Among the most abundant pathways for klebsiella are those involved in peptidoglycan maturation, glycolysis, L-1,2-propanediol degradation, myo-,chiro-and scillo-insolitol degradation, L-methionine biosynthesis and fatty acids oxidation. In addition, pathways involved in sucrose degradation; hexitol fermentation to lactate, formate, ethanol and acetate; amino acids biosynthesis; synthesis of UMP and nitrate reduction; pyruvate fermentation to isobutanol; and others, were found in klebsiella. The biosynthesis of L-alanine was predominantly attributed to klebsiella, followed by Enterobacter, *Raoultella* and *Oxalobacter*. Similarly, L-phenylalanine pathway is more abundant in klebsiella, followed by *Acinetobacter* and *Aeromonas*.

Metagenome annotation by Prokka predicted a total of 6921 different proteins, excluding the hypotheticals. Some of them are related to catabolism, anabolism, and transport of nutrients and biomolecules, regulation, DNA repair, nitrogen fixation, virulence and antibiotic resistance, among others.

Coding sequences of nitrogen-fixing genes from different species were blast to the metagenome assembly using tblastn. Non-virus, archaea or protist genes related to nitrogen-fixation were found in the microbiome of *G. berlandieri*. According to the tblastn analysis this property may be attributed to genes belonging to bacteria, most of them from the genera *Klebsiella*. In addition, the twenty genes of the Nif complex in *Klebsiella variicola* were found in the metagenome assembly by blastn (Table 2).

**Table 2.**
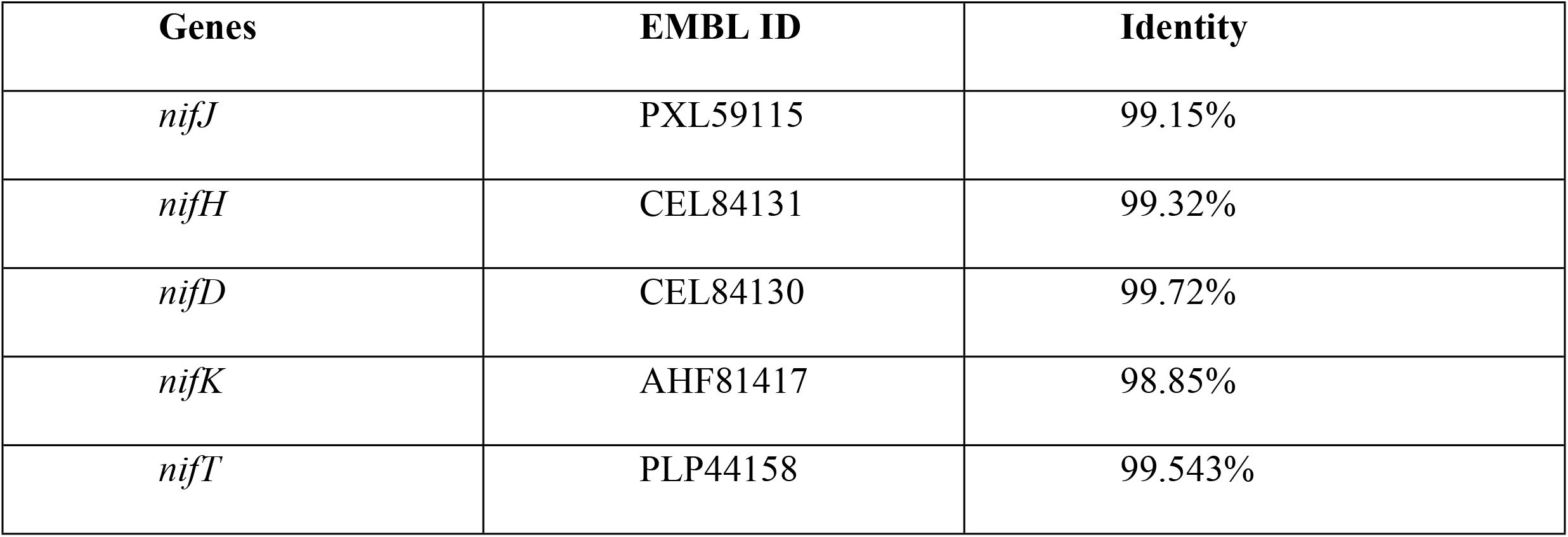
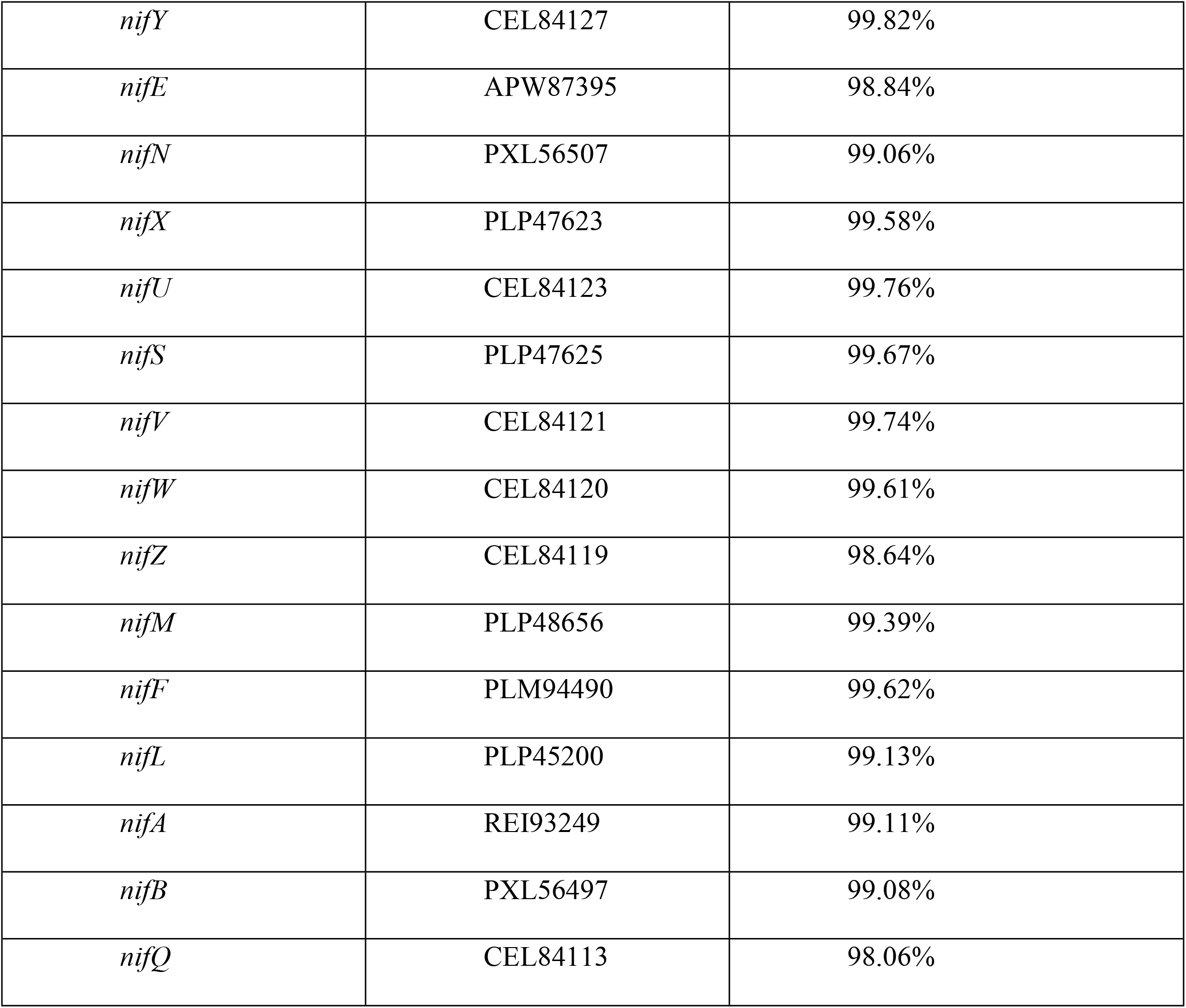
Nif genes of *Klebsiella variicola* found in *G. berlandieri* gut metagenome by blastn.

A total of 174 genes associated to virulence in Klebsiella were aligned with the metagenome assembly using blastn (29–35). According to it, 38 putative virulence genes were found in the metagenome assembly using as selection criteria an identity up to 95% throughout all the gene sequence (Table 3), of which 22 correspond to *K. variicola*, 15 belong to *K. pneumoniae* and 1 to *K. aerogenes*. The complete list of studied genes is presented as S3 Table.

**Table 3.**
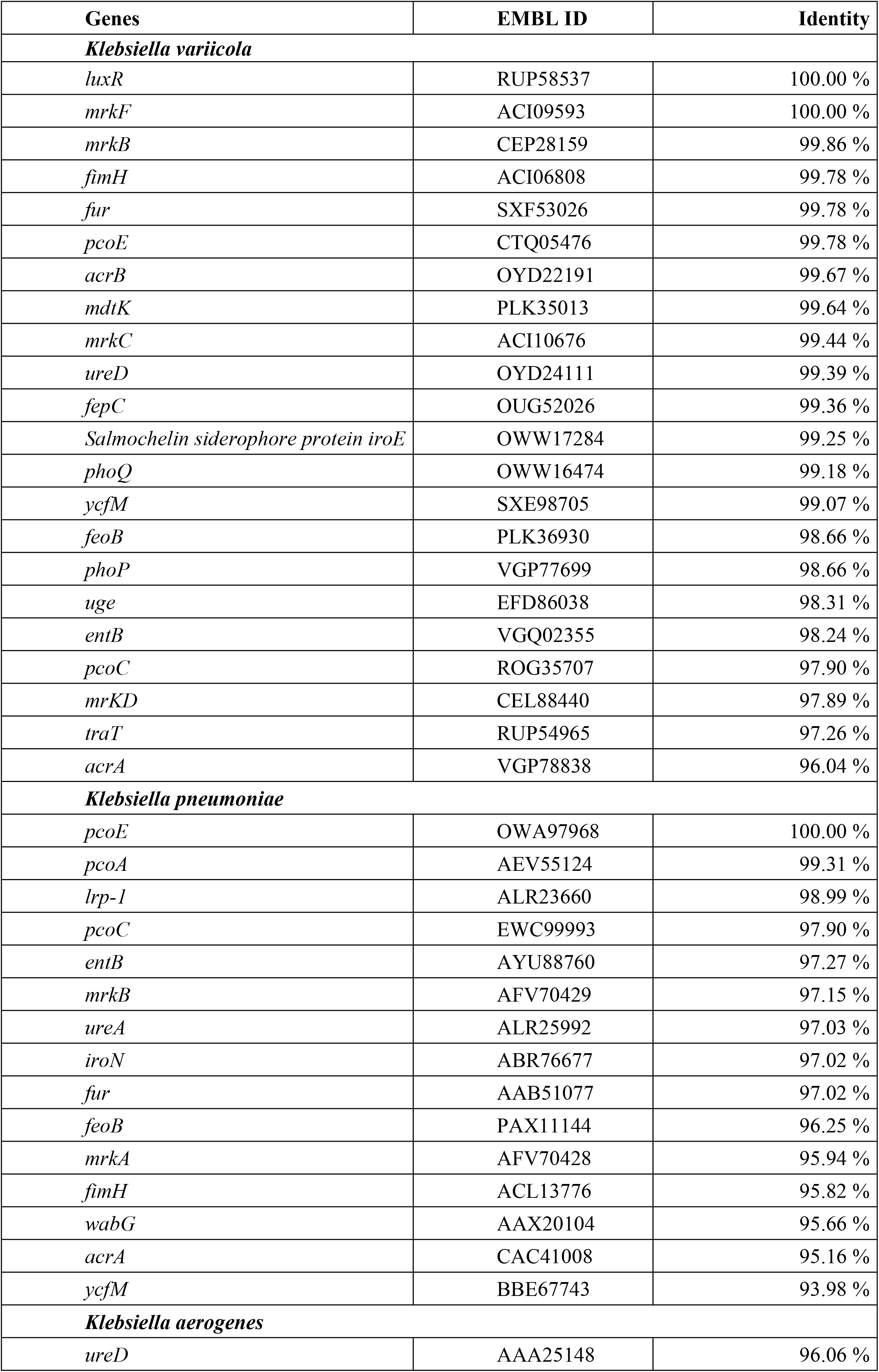
Virulence associated genes of *Klebsiella* found in *G. berlandieri* gut metagenome by blastn.

### Exo-metabolites identification

A total of 31 molecules were identified in the supernatant cultures of the *G. berlandieri* nitrogen-fixing strain KGB_5, *K. variicola* F2R9 growth separately under a microaerobic environment, in the presence and absent of a nitrogen source (S4 Table). Here, we found alanine as an exo-metabolite of KGB_5 in both nitrogen-containing and nitrogen-free medium by 1H-NMR, as well as in the extracellular culture medium of *K. variicola* F2R9 in nitrogen-containing minimal medium. A trace amount of alanine was detected in the nitrogen-containing minimal medium, with a concentration more than 30 times lower than the found in bacterial cultures grown in this medium.

Under nitrogen-free conditions, a similar metabolic footprinting was observed by ^1^H-NMR from KGB_5 and *K. variicola* F2R9. Some of the molecules produced by both strains were: 2-hydroxyisovalerate, 2-oxoglutarate, 4-hydroxyphenylacetate, betaine, fumarate, phenylacetate, pyruvate, 3-phenyllactate. However, alanine was solely detected in the extracellular culture of KGB_5; and malonate in *K. variicola* F2R9 (Fig 5).

**Fig 5.**
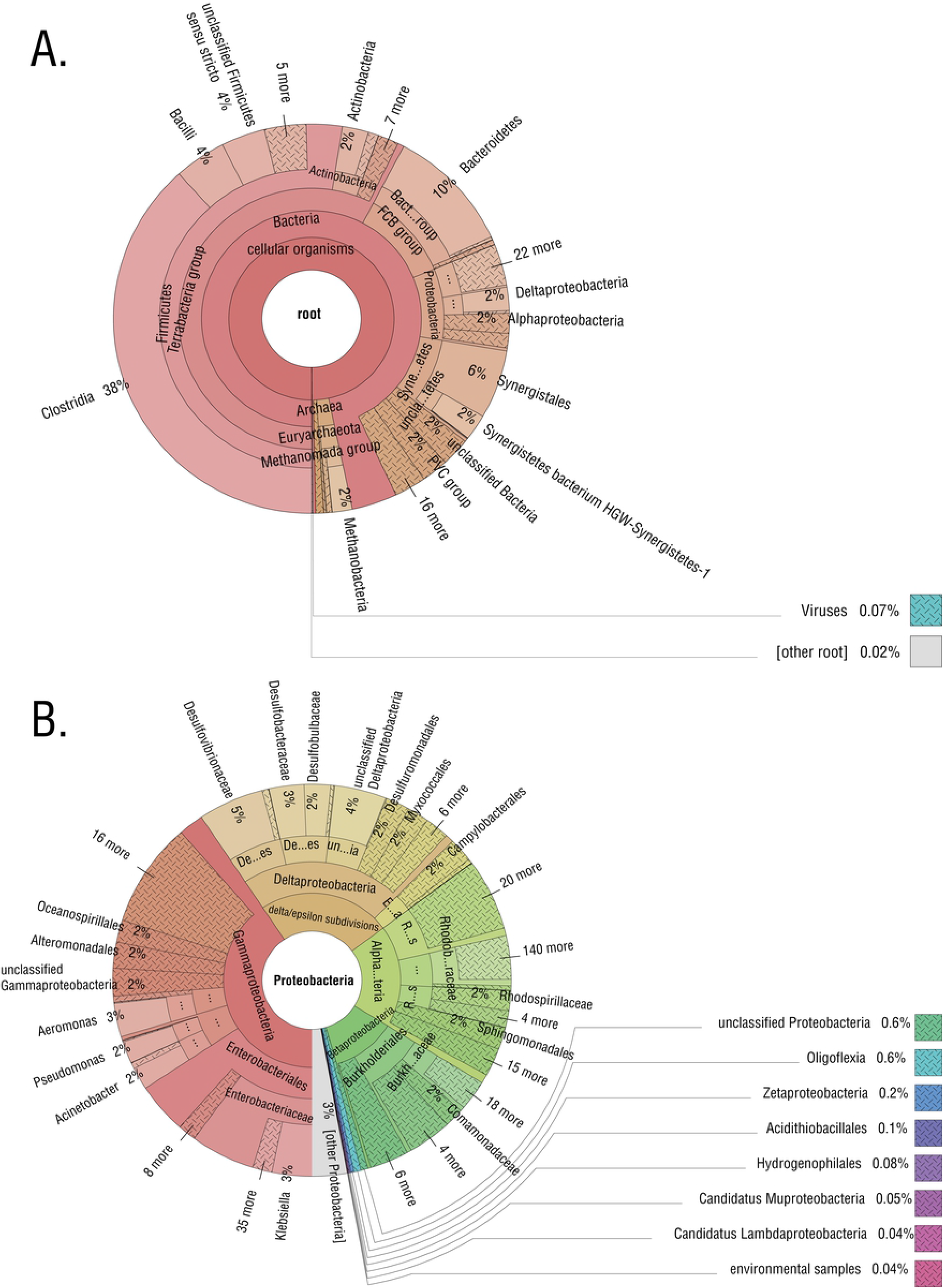
^1^H-NMR spectra of culture supernatants of A) bacterial isolated KGB_5, B) *K. variicola* F2R9 grown in C) minimal medium free of nitrogen. 1: formate; 2: phenylacetate; 3: 3-phenyllactate; 4: 4-hydroxyphenylacetate; 5: fumarate; 6: sucrose; 8: glucose; 9: 1,3-dihydroxyacetone; 10: lactate; 12: glycolate; 13: betaine; 15: methanol; 16: malonate; 17: dimethyl sulfone; 19: 2-oxoglutarate; 20: N,N-dimethylglycine; 21: dimethylamine; 22: succinate; 23: pyruvate; 24: acetone; 25: acetate; 26: alanine; 28: 3-hydroxybutyrate; 29: ethanol; 30: 2-hydroxyisovalerate and 31: 2-hydroxyvalerate.

Similarly, in nitrogen-containing minimal medium the following compounds were released by both the isolated KGB_5 and *K. variicola* F2R9: 1,3-diaminopropane, 2-hydroxyglutarate, 2-hydroxyisovalerate, 4-hydroxyphenylacetate, alanine, betaine and pyruvate. The molecules that were only detected from KGB_5 under this condition were: 3-hydroxybutyrate, fumarate, 2-hydroxyisobutyrate, glycerate and maltose. On the other hand, phenylacetate and 2-hydroxyvalerate were detected in the extracellular medium of *K. variicola* F2R9 grown in nitrogen-containing minimal medium (Fig 6).

**Fig 6.**
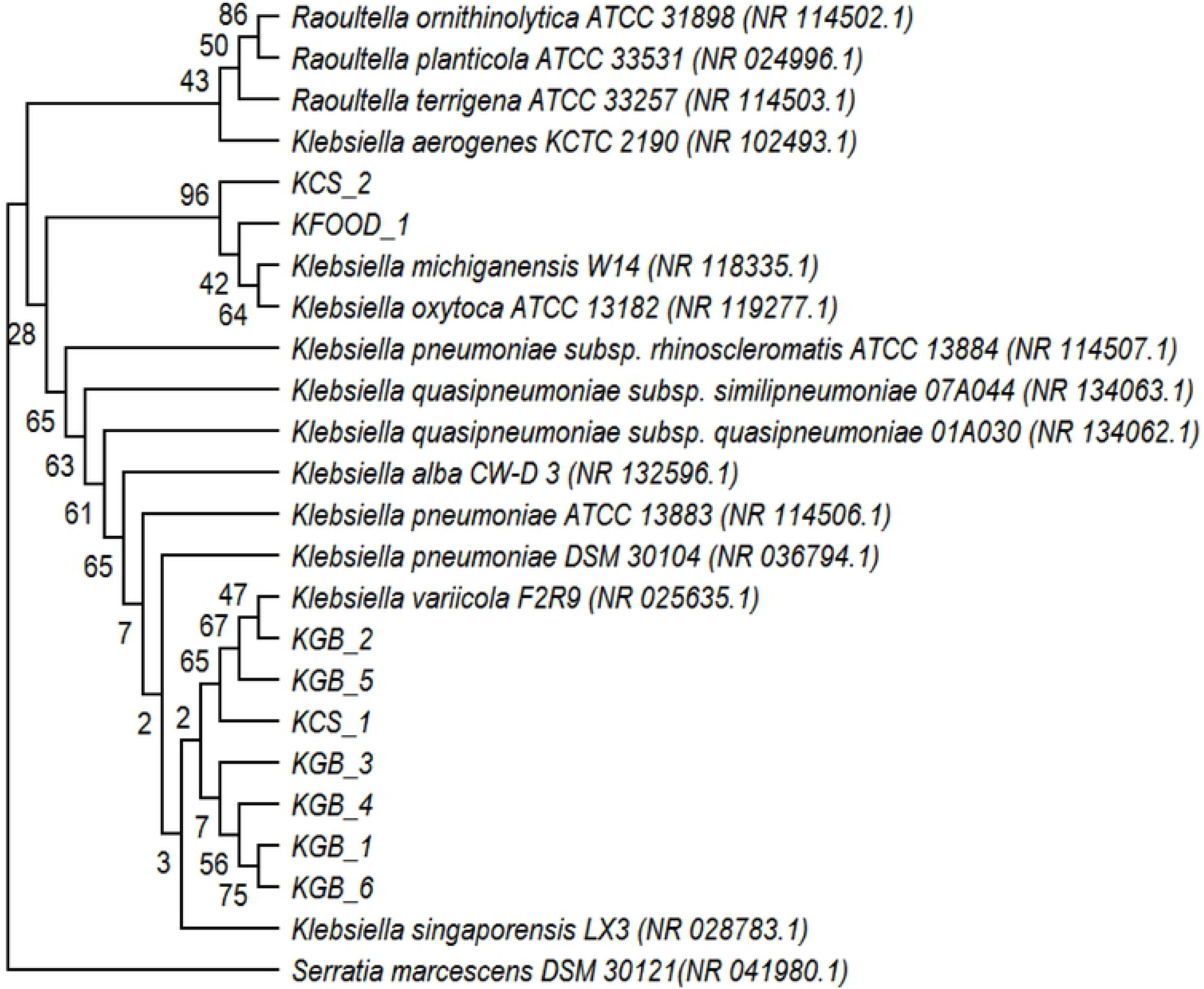
^1^H-NMR spectra of culture supernatants of A) bacterial isolated KGB_5, B) *K. variicola* F2R9 grown in C) nitrogen-containing minimal medium. 1: formate; 2: phenylacetate; 4: 4-hydroxyphenylacetate; 5: fumarate; 6:sucrose; 7: maltose; 8:glucose; 9: 1,3-dihydroxyacetone; 10: lactate; 11: 2-hydroxyglutarate; 12: glycolate; 13: betaine; 14: glycerate; 15: methanol; 16: malonate; 17: dimethyl sulfone; 18: 1,3-diaminopropane; 20: N,N-dimethylglycine; 21: dimethylamine; 22: succinate; 23: pyruvate; 24: acetone; 25: acetate; 26: alanine; 27: 2-hydroxyisobutyrate; 28: 3-hydroxybutyrate; 29: ethanol; 30: 2-hydroxyisovalerate and 31: 2-hydroxyvalerate.

## Discussion

Bacterial diversity from feces has been extensively studied in animals including humans with different diets, however the microbiota from tortoises had not been reported. Insects that feed on low nitrogen and high carbon diets profit from nitrogen-fixing symbioses with distinct gut bacteria. Termites are the best known examples, but wood eating beetles also harbor diazotrophs. Nitrogen-fixing symbioses had not been described in reptiles and may help to compensate nitrogen deficiencies in their diet.

### Nitrogen-fixing bacteria in tortoise feces

Culture-dependent and independent approaches revealed that the enterobacteria *K. variicola* is found in *G. berlandieri*. From *C. sulcata*, *K. variicola* was isolated from feces in culture media as well. Here fecal bacterial isolation was in microaerobic nitrogen-free medium to enrich for nitrogen-fixing bacteria. This allowed the isolation of nitrogen-fixing but few non-nitrogen-fixing bacteria were obtained as well that were identified as Rhanella.

The culture independent approach using metagenomics suggested that *K. variicola* may be responsible for nitrogen fixation in *G. berlandieri* guts as the only *nif* genes recovered corresponded to *K.variicola*.

*K. variicola* is a nitrogen-fixing species closely related to *K. pneumoniae* (36, 37) which rarely fixes nitrogen. The former has been isolated from plants such as maize, sugarcane and banana, as a plant-growth-promoting root endophyte (inside plant tissues). However, some *K. variicola* strains have been also isolated from hospitalized patients, in special from babies. Klebsiellas are common colonizers of human intestine. From human stools, klebsiellas have been isolated even from babies feeding from breast-milk (38). Notably the use of *K variicola* in agricultural crops to promote plant growth has been dismissed due to their pathogenic potential (39). However it is unlikely that *K. variicola* is pathogenic in tortoises as they were healthy and we suppose that they are mutualists as occurs in plants. Here we surmise that nitrogen fixation indeed occurs in tortoise guts on the basis of the ethylene reduction activity that we detected in the excreted feces. Therefore our hypothesis is that nitrogen-fixing bacteria may benefit the host by supplying nitrogen that seems to be deficient in the tortoise diet.

Environmental *K. variicola* strains generally have fewer antibiotic resistance genes than clinical isolates (36), here we found that klebsiellas from feces were susceptible to Cb100 and Nal20, though they were resistant to ampicillin. All *K. variicola* strains are resistant to ampicillin and the gene responsible is chromosomally encoded indicating that is not easily lost or transferred. Previously we proposed that there exists a different mode of transmission for *K. variicola* and *K. pneumoniae* with *K. variicola* acquired from the environment and *K. pneumoniae* transmitted from human to human (36, 40). If tortoise klebsiellas are transmitted from animal to animal, then better gut adapted strains (see below) may be selected avoiding infection with pathogenic bacteria. Indeed we found that *K. variicola* from tortoises had a low proportion of virulence determinants.

### Origin of tortoise klebsiellas

Bacteria may be obtained from feces ingestion as coprophagy is a common practice among some tortoises or they may derive from their vegetable diet as *K. variicola* strains are commonly found in plants. To address this we isolated bacteria associated from the vegetables used for feeding the Texas tortoises or the pellets used to feed the African tortoise. All plants contain endophytes and there is a large diversity of endophytes which may be recovered from plants after a surface disinfection procedure that eliminates bacteria that are not within tissues. We suppose that by being endophytes, klebsiellas are protected in the inside of vegetable tissues and may survive the digestion process. Since herbivorous animals exhibit the largest diversity of gut bacteria in comparison to ommivorous or carnivorous animals (41), it seems that endophytes may become constituents of gut microbiota, at least transiently. Alternatively, common klebsiella symbionts among tortoises would indicate that they are vertically transmitted and well adapted to hosts. We found that diet and gut nitrogen-fixing bacteria were not the same, indicating that there is not a direct transfer of nitrogen-fixing bacteria from the ingested food. If klebsiellas are not obtained from the diet, then gut klebsiellas may be obtained by coprophagy in tortoises and perhaps early in life. It is remarkable that fecal klebsiella from distinct individual tortoises are very similar from the REP-BOX study performed (that allows strain distinction), the limited diversity encountered would favor the tortoise-tortoise transmission hypothesis.

It is worth considering that there could be some negative aspects of nitrogen fixation by *Klebsiella* in guts. Antibodies against nitrogenase have been detected in humans with ankylosing spondylitis and *Klebsiella* infections have been considered as candidates for the onset of such disease (42). We wonder if tortoises would have as well an immune response towards klebsiella nitrogenase and what the effects in their heath could be.

Reptile gut bacteria include clostridia as important components (43), as we found in the metagenome analysis of *G. berlandieri*.

### Exo-metabolite analysis

Diazotrophs such as *Klebsiella* that are among the free-living nitrogen-fixing bacteria normally do not excrete ammonium produced by nitrogen fixation and use it for their own bacterial growth. To further explore this we analyzed the metabolites excreted by a *K. variicola* isolate *in vitro* under microaerobic conditions in nitrogen free medium in comparison to medium with added nitrogen. By 1H-NMR exo-metabolite analysis, alanine was detected in the supernatant of *K. variicola* KGB_5 grown in nitrogen free medium and microaerobic conditions while the plant *K. variicola* F2R9 strain did not excrete alanine under those conditions. Interestingly some *Rhizobium* strains may provide alanine to plants in symbiosis (44, 45). It would be worth exploring if alanine excretion is a common adaptative characteristic of tortoise klebsiellas and not particular to the tested strain. In addition, a number of exo-metabolites were produced by both KGB_5 and *K. variicola* F2R9 under nitrogen-containing minimal medium and nitrogen free conditions.

## Acknowledgements

To Michael Dunn for reading the manuscript.

## Funding Sources

This research was supported by Universidad Autónoma de México [Grant: Programa de Becas Posdoctorales en la UNAM 2016 to D. M-G], CONACyT [Grant: 253116] and PAPIIT [Grant: IN207718]. This study made use of UNAM’s NMR lab: LURMN at IQ-UNAM, which is funded by CONACYT Mexico (Project: 0224747), and UNAM.

## Supporting information captions

**S1 Table. Taxonomic classification of the metagenomic reads from the *G. berlandieri* gut microbiome.**

**S2 Table. HUMAnN2 functional analysis results**.

**S3 Table. List of genes associated to virulence in *Klebsiella* used in this study.**

**S4 Table. Chemical shifts of the compounds identified by 1H NMR (700 MHz, 298K, TSP=1mM, D2O sodium phosphate buffer 0.123M at pH 7.4).** TSP was used as internal standard and referenced to chemical shift 0.000 ppm.

